# Cognitive Deficits and Altered Cholinergic Innervation in Young Adult Mice Carrying a Parkinson’s Disease LRRK2-G2019S Knockin Mutation

**DOI:** 10.1101/2022.01.26.477929

**Authors:** Ayan Hussein, Alexander Tielemans, Mark G. Baxter, Deanna L. Benson, George W. Huntley

## Abstract

Impaired executive function is a common and debilitating non-motor symptom of idiopathic and hereditary Parkinson’s disease (PD), but there is little understanding of the underlying pathophysiological mechanisms and circuits. The G2019S mutation in the kinase domain of leucine-rich repeat kinase 2 (*LRRK2*) greatly increases risk for late-onset PD, and non-manifesting *LRRK2*-G2019S carriers also exhibit early and significant cognitive impairment. Here, we subjected young adult mice carrying a *Lrrk2*-G2019S knockin mutation to touchscreen-based operant tasks that measure attention, goal-directed learning and cognitive flexibility, all of which rely on prefrontal-striatal connectivity and are strongly modulated by cholinergic innervation. In a visuospatial attention task, mutant mice exhibited significantly more omissions and longer response latencies than controls that could not be attributed to deficits in motivation, visual sensory perception per se or locomotion, thereby suggesting impairment in divided attention and slower information processing speed. Pretreating mice with the acetylcholinesterase inhibitor donepezil normalized both higher omission rates and longer reward latencies in the mutants, but did not affect any performance metric in controls. Strikingly, cholinergic fiber density in mPFC and dorsomedial striatum was significantly sparser in mutants than in controls, while further behavioral interrogation of the mutants revealed significant impairments in action-outcome associations but preserved cognitive flexibility. These data suggest that the G2019S mutation impacts cholinergic innervation and impairs corticostriatal network function in young adulthood that may contribute to early PD-associated cognitive deficits.

**STATEMENT OF SIGNIFICANCE:** The *LRRK2*-G2019S mutation causes hereditary Parkinson’s disease and is found in some idiopathic cases. Early cognitive impairment is a common symptom of hereditary and idiopathic PD, yet there is little mechanistic understanding of such impairment. Here, we tested young adult *Lrrk2*-G2019S knockin mice in a series of touchscreen-based visuospatial tasks. We found that mutants exhibited significant deficits in attention and goal-directed learning, and had significantly slower information processing speed. Treatment with an acetylcholinesterase inhibitor reversed some of these behavioral deficits, while anatomical analyses showed significantly sparser cholinergic innervation of brain structures important for executive function. These findings suggest the G2019S mutation alters cholinergic signaling in young adulthood, and thus may contribute to early PD-associated impairment in several cognitive domains.

## INTRODUCTION

Cognitive impairment affects 30-50% of those with Parkinson’s disease (PD), with a greater prevalence in males and an onset that can precede motor symptoms (Muslimovic et al., 2005; Cammisuli et al., 2019). Deficits include impaired attention, cognitive flexibility, perceptual decision making, reinforcement learning and generally slower information processing speed, and can progressively worsen over time leading to dementia (Robbins and Cools, 2014; Perugini et al., 2018). The negative impact of PD-associated cognitive impairment on quality of life is significant, often exceeding that resulting from motor disturbances (Prakash et al., 2016). Nevertheless, there is little understanding of pathophysiological mechanisms and circuits that underlie PD-associated cognitive impairment, hindering treatment options.

In primates and rodents, executive function requires prefrontal cortex and connectivity with striatum (Selemon and Goldman-Rakic, 1988; Yin et al., 2005; Ragozzino, 2007). Humans with early PD-associated impairment in executive function show altered frontal-striatal network activity (Lewis et al., 2003; Wolters et al., 2019). Additionally, executive function, particularly attention, is strongly modulated by cholinergic innervation of prefrontal cortex and striatum (Everitt and Robbins, 1997; Baxter and Chiba, 1999; Ballinger et al., 2016). Cholinergic innervation of prefrontal cortex arises from neurons in basal forebrain and from cortical cholinergic interneurons (Ballinger et al., 2016; Obermayer et al., 2019), while cholinergic innervation of striatum arises largely from striatal cholinergic interneurons with a sparse contribution from neurons in mesopontine tegmental nuclei (Bolam et al., 1984; Dautan et al., 2016). While much focus of PD research is dopamine neuron degeneration, cholinergic neurons also degenerate in humans during the course of PD (Choi et al., 2012; Pasquini et al., 2021). Progressive loss of cortical cholinergic projections is associated with cognitive impairment in non-demented PD patients (Bohnen et al., 2015; Kim et al., 2017) and has been associated with PD-related deficits in attention and executive function (Bohnen et al., 2006). Thus, while dysfunctional cholinergic signaling is implicated in PD-related cognitive decline, there is little understanding of how, when or where changes in cholinergic signaling arise.

One approach to address such lapses is to experimentally interrogate animals carrying genetic mutations that in humans cause PD (Dawson et al., 2010). The G2019S autosomal dominant mutation in leucine-rich repeat kinase 2 (*LRRK2*) is the most frequent cause of familial PD and is also found in some idiopathic PD cases (Martin et al., 2014). LRRK2 is a large protein with a catalytic core comprising GTPase and kinase domains. The G2019S mutation, located in the kinase domain, increases kinase activity (Paisan-Ruiz et al., 2013). Human G2019S carriers develop late-onset PD with a clinical presentation similar to idiopathic PD, suggesting common pathophysiological mechanisms (Martin et al., 2014). Consistent with this, G2019S carriers also exhibit early cognitive impairment, although this tends to be less severe than idiopathic PD (Healy et al., 2008; Thaler et al., 2012; Gaig et al., 2014; Mirelman et al., 2015; Ben Romdhan et al., 2018; Piredda et al., 2020) and, prior to the onset of motor symptoms, exhibit altered functional coupling between cortex and striatum (Thaler et al., 2013; Helmich et al., 2015; Bregman et al., 2017). Additionally, human G2019S carriers with or without PD exhibit altered acetylcholinesterase activity in cortical and subcortical regions in comparison with idiopathic PD or healthy controls (Liu et al., 2018).

In human and rodent brains, LRRK2 expression is highest in striatum and cortex, with low expression in substantia nigra pars compacta or basal forebrain (Benson and Huntley, 2019). Developmental studies of postnatal mice show that LRRK2 expression levels in cortex and striatum rise contemporaneously with corticostriatal synaptogenesis (Giesert et al., 2013), suggesting that the mutation may shape development of circuits critical for executive function in ways that negatively impact cognitive abilities later in life (Hussein et al., 2021). Accordingly, in this study, we use G2019S knockin mice to test the prediction that executive function across several cognitive domains is aberrant by young adulthood, and explore functional and anatomical aberrations in cholinergic innervation as a contributing factor, both of which have not previously been rigorously explored in PD mouse models.

## MATERIALS AND METHODS

### Mice

PD is more common in males than in females and at the time these studies were initiated, most evidence supported that early cognitive decline in idiopathic PD may be worse in males than in females (Elbaz et al., 2016; Reekes et al., 2020). Accordingly, the present study used male *Lrrk2*-G2019S knockin (GSKI) mice (RRID:MGI:6273311) that were congenic on a C57Bl/6NTac background. It is worth noting, however, that some newer work suggests that within PD patients, cognitive impairment in male *LRRK2*-G2019S carriers may be less severe than in female carriers (Ben Romdhan et al., 2018). In any event, over the course of all experiments, mice were aged 2-6 postnatal months. Age- and strain-matched wildtype (WT) mice served as controls. All mice were bred and raised under identical conditions in the vivarium. The GSKI mice were generated by Eli Lily and characterized previously (Matikainen-Ankney et al., 2016, 2018; Guevara et al., 2020). They exhibit normal life spans and body weights, express normal levels of tyrosine hydroxylase in the striatum, and express normal physiological levels of LRRK2-G2019S protein in brain. GSKI mice were backcrossed to commercial WT C57Bl/6NTac mice every fourth generation to prevent genetic drift. Genotypes were confirmed by PCR and sequencing. In humans, the G2019S mutation is autosomal dominant, producing similar disease onset and progression in heterozygous and homozygous mutation carriers (Zimprich et al., 2004; Paisán-Ruiz et al., 2013). As predicted from this, previous studies showed similar behavioral and electrophysiological outcomes in heterozygous and homozygous G2019S mice (Matikainen-Ankney et al., 2016, 2018; Guevara et al., 2020). We thus used heterozygous G2019S mutants for these experiments. All procedures were approved by Mount Sinai’s Institutional Animal Care and Use Committee and conformed to guidelines established by the National Institutes of Health.

### Behavioral tasks

Mice were 2-to 3-months old when behavioral experiments began. They were housed in groups of 3-4 in a temperature-controlled room with a 12 hr light/dark cycle (lights on at 7:00 am). Behavioral testing was conducted once a day for 5-6 days a week during the light phase of the cycle. Mice were food restricted to 85% of their free-feeding weight. Water was available *ad libitum*, and food rations, adjusted based on body weight, were given at the end of each day.

### Apparatus

All behavioral tasks were carried out using four automated Bussey-Saksida style touchscreen-based operant chambers (Lafayette Instruments, Lafayette, IL) consisting of four modular testing chambers housed individually within a sound- and light-attenuating cubicle equipped with an overhead camera to monitor task-related locomotion. Inside each chamber, a reward collection magazine connected to a liquid dispenser pump was situated opposite a touch-sensitive monitor. Depending on the task, the touchscreen monitor was divided into one, two or five response windows by interchanging different black Perspex masks. Each response window was 4.0 x 4.0 cm regardless of how many response windows each mask contained. The collection magazine contained an LED illuminated coincident with delivery of a liquid reward (Nesquik^®^ strawberry-flavored milk) and a tone. Infrared beam arrays distributed in front of the touchscreen, reward collection port and across the intervening floor, were used to monitor locomotor. Different training schedules, depending on the task, were designed and data were analyzed using ABET II Touch software (Lafayette Instruments), while inputs and outputs from all four chambers were controlled by WhiskerServer software (Lafayette Instruments).

### 5-CSRT task

This task was conducted on 18 GSKI mice and 19 WT mice and followed established procedures (Bari et al., 2008; Mar et al., 2013). The following performance metrics were collected (and defined as): % response accuracy (# correct responses/(total number of correct and incorrect responses) x 100); % correct responses (# correct responses/(total number of correct, incorrect and omitted responses) x 100); % omissions (# of omissions/(total number of correct, incorrect and omitted responses) x 100); # trials (# correct and incorrect trials); correct response latency (the elapsed time between stimulus offset following a correct touch and head entry into the reward magazine signaled by a back IR beam break).

### Pre-training

Mice progressed through 5 pretraining stages (habituation, initial touch, must touch, must initiate and punish incorrect) before advancing to training. In stage 1, they were acclimated to the chambers for a 40 min session with the tray-light turned on, and the tray was primed with strawberry milk. In stages 2 and 3, a mask with five response windows was placed over the touchscreen, and mice learned to nose poke the stimulus (white square) displayed randomly in one of the five response windows to elicit strawberry milk delivery. In stage 4, mice learned to initiate each trial via head entry into the magazine tray; withdrawal from the magazine elicited the next stimulus presentation. Mice were required to complete 30 trials in 60 minutes across two consecutive days to meet completion criteria for stages 2-4. In stage 5, mice were discouraged from nose-poking blank locations on the screen by imposing a 5 sec timeout period signaled by illumination of the house light. Mice were rewarded with 7 μl of strawberry milk at all stages. To progress to task training, mice were required to complete 24 out of 30 trials in 60 minutes (80% accuracy) across two out of three consecutive days.

### Training

Daily training sessions lasted 60 min. At the beginning of each session, the magazine light was illuminated and a free strawberry milk reward was dispensed. A nose poke entry into the reward port extinguished the magazine light, and upon exiting initiated a 5 sec intertrial interval (ITI). Following the ITI, a visual stimulus (white square) was randomly displayed in 1 of 5 locations on the screen. A nose poke response made in the corresponding location either during the stimulus presentation or the following 5 sec limited hold period (stimulus offset) was deemed correct and resulted in strawberry milk delivery. The stimulus duration was initially set at 32 seconds and gradually reduced to 16, 8, 4, and 2 sec to establish baseline performance at 2 sec stimulus duration. The performance criteria were completion of 50 out of 60 trials within 60 min, response accuracy ≥80%, and trial omissions ≤20%, across three of four consecutive days.

### Probe trials

Once all mice were performing at baseline (2 sec stimulus duration), they were subjected to two probe trials in which attentional demand was increased by shortening stimulus durations. In probe trial 1 (n=19 WT mice, 17 GSKI mice), stimulus durations were (in seconds): 1.5, 1.0, 0.8, 0.6, and 0.4, where mice were tested at a single stimulus duration for two consecutive days (within session). The order of stimulus durations presented to each mouse was randomized in a Latin square design. In between each of the shortened stimulus durations, mice were tested at the baseline 2 sec stimulus duration until performance criteria were reattained (completion of 50 out of 60 trials, ≥80% accuracy and ≤20% omissions in 60 min). In probe trial 2 (n=13 WT mice, 11 GSKI mice), stimulus durations were (in seconds): 2.0, 1.0, 0.8 and 0.6 in a within-session block design for two consecutive days. Equal numbers of each of the four stimulus durations were randomly presented during the 60-trial session over a 60 min period.

### Visual Perception test

We tested visual perceptual abilities of mice at the shortest stimulus duration used in the 5-CSRT task (0.4 sec) in a subset of mice completing the 5-CSRT task (n = 13 WT mice; 9 GSKI mice), under conditions where attentional demand was lowered by presenting the stimulus in one response window, thereby eliminating the need for mice to divide their attention across 5 response windows. In this 1-choice version of the task, the stimulus (white square) was only presented in the middle window of the five-response window mask. Other task parameters and the primary response rules remained the same. Once mice reattained the baseline performance at 2 sec, they were tested on the 0.4 sec stimulus duration (50 out of 60 trials within 60 min) for two consecutive days.

### Administration of donepezil

These experiments were conducted on a subset of WT (n=13) and GSKI (n=9) mice after completing the two probe trials. Mice received either donepezil or vehicle (normal saline) using a within-subject design where, for each mouse, the order of treatment with vehicle or donepezil was counterbalanced across animals of each group. Donepezil hydrochloride (0.3 mg/kg; Bio-Techne Corp, Minneapolis, MN) was dissolved in 0.9% saline and was made fresh each day. The donepezil dose used was based on prior studies (Yuede et al., 2007; Bartko et al., 2011; Romberg et al., 2011). Mice were injected intraperitoneally one hour before behavioral testing on each day of a two-day testing sequence in the 5-CSRT task. Testing consisted of 50 trials for 2 consecutive days at a stimulus duration of 0.4 sec. In between dosing/testing sequences, mice received three drug-free days over which they were tested using a baseline 2 sec stimulus duration; animals were required to reinstate baseline performance (≥80% accuracy, <20% omissions).

### Instrumental conditioning and outcome devaluation

Action-outcome learning was assessed in a second, separate cohort of mice (n=7 WT and 6 GSKI mice) by adapting for touchscreens a previously described 4-day lever-pressing paradigm of instrumental conditioning followed by subsequent tests of reward devaluation to discriminate goal-directed and habitual responding (Shan et al., 2014). In this task, a one response-window mask was used. Training consisted of two 60-minute sessions of habituation in the chamber as described above, followed by a 2 hr session using a continuous reinforcement (CRF) schedule, where each nose poke to the stimulus window elicited a reward. Following training, mice progressed to a 4-day testing protocol, where each session was limited to 60 min. On day 1, mice were tested on a CRF schedule. On days 2-4, mice were subjected to a random interval (RI) session of 15, 30, and 30 sec. On days 5 and 6, outcome devaluation tests were conducted. On day 5, half of the mice were given 1 hr free access to strawberry milk reward (devalued condition), while the other half got 1 hr free access to home-cage chow (non-devalued condition), followed immediately by a 5 min extinction test. No reward was elicited in this extinction test. On day 6, a second extinction test was performed in the same manner as the firstexcept that the satiety conditions were reversed.

### Pairwise visual discrimination and reversal learning

Cognitive flexibility was tested in a third, separate cohort (n=5 WT and 7 GSKI mice) using pairwise visual discrimination and reversal learning. Before testing, mice underwent touchscreen training as described above, but using a mask containing two response windows. In the initial discrimination phase, mice were presented simultaneously with two distinct visual stimuli (plane vs. spider) and taught to associate one stimulus (e.g. plane) with strawberry milk delivery and the other (e.g. spider) with a 5 sec timeout period signaled by illumination of the house light (Bartko et al., 2011). The rewarded stimulus was pseudo-randomly displayed in one of the two response windows and was counterbalanced within mice. An incorrect choice resulted in a correction trial in which the stimulus display was repeated in the same window until the mouse made a correct response. However, correction trials were not included in the total tally of correct trials. Upon completing 24 out of 30 trials (80% accuracy) within 60 mins in two consecutive days, mice advanced to the subsequent reversal phase in which reward contingencies were switched. As with the discrimination phase, an incorrect choice resulted in a correction trial in which the stimulus display was repeated in the same window until the mouse made a correct response. Measurements included the rate at which mice acquired the new stimulus-reward associations as well as the number of correction responses in each phase (Bussey et al., 1997).

### Progressive ratio task

Motivation for the reward was evaluated by a progressive ratio test of subsets of mice following completion of the 5-CSRT task (n=13 WT and 11 GSKI mice) or instrumental conditioning/outcome devaluation (n=4 WT and 4 GSKI mice). Mice were subjected to fixed ratio (FR) schedules in which nose-poking the illuminated center response window for a fixed number of times (FR 1, 2, 3, or 5) was required to elicit a strawberry milk reward. Criterion to advance was completion of 30 trials within 60 min for two consecutive days. Following completion of FR5, mice advanced to a progressive ratio (PR) 4 testing schedule, where reward response requirement incremented on a linear + 4 basis (i.e., 1, 5, 9, 13, etc.) in each subsequent trial. Sessions were run for 60 minutes and were terminated when a mouse ceased responding for 5 minutes (the breakpoint).

### Statistical analysis of behavioral data

Statistical analyses were performed in R version 4.1.2 (R Core Team, 2021). Statistical tests for each dataset are specified within each figure legend. For most analyses we used a linear mixed models regression framework with genotype and behavioral variables (e.g. stimulus duration) or treatment levels (donepezil/saline) as fixed effects and mouse as a random effect, using R packages lme4 (Bates et al., 2015) and lmerTest (Kuznetsova et al., 2017) and the mean score in each session as the unit of analysis. These models were evaluated with F-tests on type III sums of squares on fixed effects and their interactions. Where significant effects were indicated, these were decomposed through comparisons of estimated marginal means using the emmeans package (Lenth, 2022). Data are reported as mean ± standard error of the mean (SEM) and all significant comparisons are displayed by asterisks in the figures.

### Immunolabeling

Behaviorally naïve male mice (n=3 WT, 3 GSKI mice, aged 2-3 months) were anesthetized with a mixture of ketamine (90 mg/kg) and xylazine (20 mg/kg, IP) and transcardially perfused first with cold 1% paraformaldehyde (PFA) in 0.1M phosphate buffered saline (PBS, pH 7.3) for 30-45 sec followed by cold 4% PFA for 10 min. Brains were dissected, post-fixed for 6-8 hours in 4% PFA, then soaked in 7% sucrose in PBS for 24 hrs. Serial coronal tissue sections through mPFC (including areas PL/IL) and dorsal striatum were cut on a vibratome at a setting of 50 μm. Tissue sections were then permeabilized with 0.25% Triton X-100 in PBS for 15 min, washed thoroughly in PBS, and incubated in a PBS solution containing 10% Bovine Serum Albumin (BSA) for 1.5 h at room temperature (RT). Sections were then incubated for 72 hrs at 4°C in primary rabbit anti-vesicular acetylcholine transporter (VAChT) antisera (Synaptic Systems; 1:500, RRID_AB887864) containing 2% BSA and 0.1% Triton X-100 in PBS. Sections were washed thoroughly in PBS, followed by RT incubation for 1 hr with secondary donkey anti-rabbit Alexa 488 (Jackson ImmunoResearch; 1:500, RRID_AB2313584). After washing, sections were mounted using VectaShield antifade mounting media and counterstained with DAPI. Coverslip edges were sealed with nail polish and dried overnight in the dark.

### Image analysis

Analysis was carried out on three immunolabeled sections through areas PL/IL and three immunolabeled sections through dorsomedial striatum from each mouse. No attempt was made to analyze PL and IL separately. A series of contiguous single optical plane images through PL/IL or striatum were acquired with a 40X objective on a Zeiss LSM 780 confocal microscope and stitched together using Zeiss Zen SP1 (black) software (v.8.1.11.484) to create a single tiled image per section through each structure (tiled image dimensions in PL/IL: 1020 μm x 1226 μm, spanning layers 1-6; tiled image dimension in DMS: 820 μm x 820 μm). Image acquisition parameters were kept identical for all sections from mice of each genotype. A sampling frame (207.5 μm^2^) was used to define a region of interest (ROI) and overlayed onto the raw tiled images of PL/IL (6 ROIs pers section, sampling superficial-to-deep cortical layers) and dorsomedial striatum (3 ROIs per section). VAchT fiber density, average size of immunolabeled puncta and immunofluorescent intensity were determined using ImageJ by applying a thresholding function to each ROI to create a binarized mask of the immunofluorescent signal. The mask was then applied to the original image to capture and quantify density, area and intensity of suprathreshold immunofluorescent pixels. The density of VAChT immunolabeled fibers was expressed as the percent of R0I area occupied by suprathreshold pixels (Dougherty et al., 2020). Statistical comparisons between groups were made using a linear mixed model on each measure, with region and genotype as fixed effects and mouse as a random effect; p values < 0.05 were considered statistically significant.

## RESULTS

Cohorts of young adult male wildtype (WT) and *Lrrk2*-G2019S knockin (GSKI) mice were subjected to visuospatial tasks that evaluate attention, goal-directed learning and cognitive flexibility using automated, standardized touch screen-based operant chambers.

### The 5-CSRT task reveals an attention deficit in GSKI mice

Task acquisition consisted of an initial training phase in which mice learned to respond to a stimulus presented on a screen for a limited time. In the training phase, stimulus duration was initially 32 sec, then sequentially reduced to a baseline of 2 sec. Performance criteria for advancing to the next shorter stimulus duration were completion of ≥50 trials within a 60 min session, with response accuracy of ≥80% and an omission rate of ≤20% across 3 out of 4 consecutive sessions (Mar et al., 2013). All WT and GSKI mice reached task criteria at each stimulus duration. However, while GSKI mice displayed similar percentages of correct responses at each stimulus duration (**Fig. 1A**), they made more omissions (**Fig. 1B**) and completed fewer trials (**Fig. 1D**) compared to WT mice. GSKI mice exhibited longer latencies compared to WT mice at the longer stimulus durations during initial training (**Fig. 1E**). During the ITI, GSKI mice made fewer premature responses compared to WT mice (**Fig. 1F**). Additionally, GSKI mice exhibited greater response accuracy (correct trials / correct + incorrect trials) in comparison with WT mice (**Fig. 1C**), perhaps reflecting a tendency to fail to respond (make an omission) versus making an incorrect response on trials where there was a lapse in attention. There were no significant differences between genotypes in perseverative responses following a correct choice at any stimulus duration (genotype, F_1,35_ = 0.64, p=0.43, stimulus duration x genotype interaction: F_4,510_ = 0.72, p=0.58).

**Figure 1.**
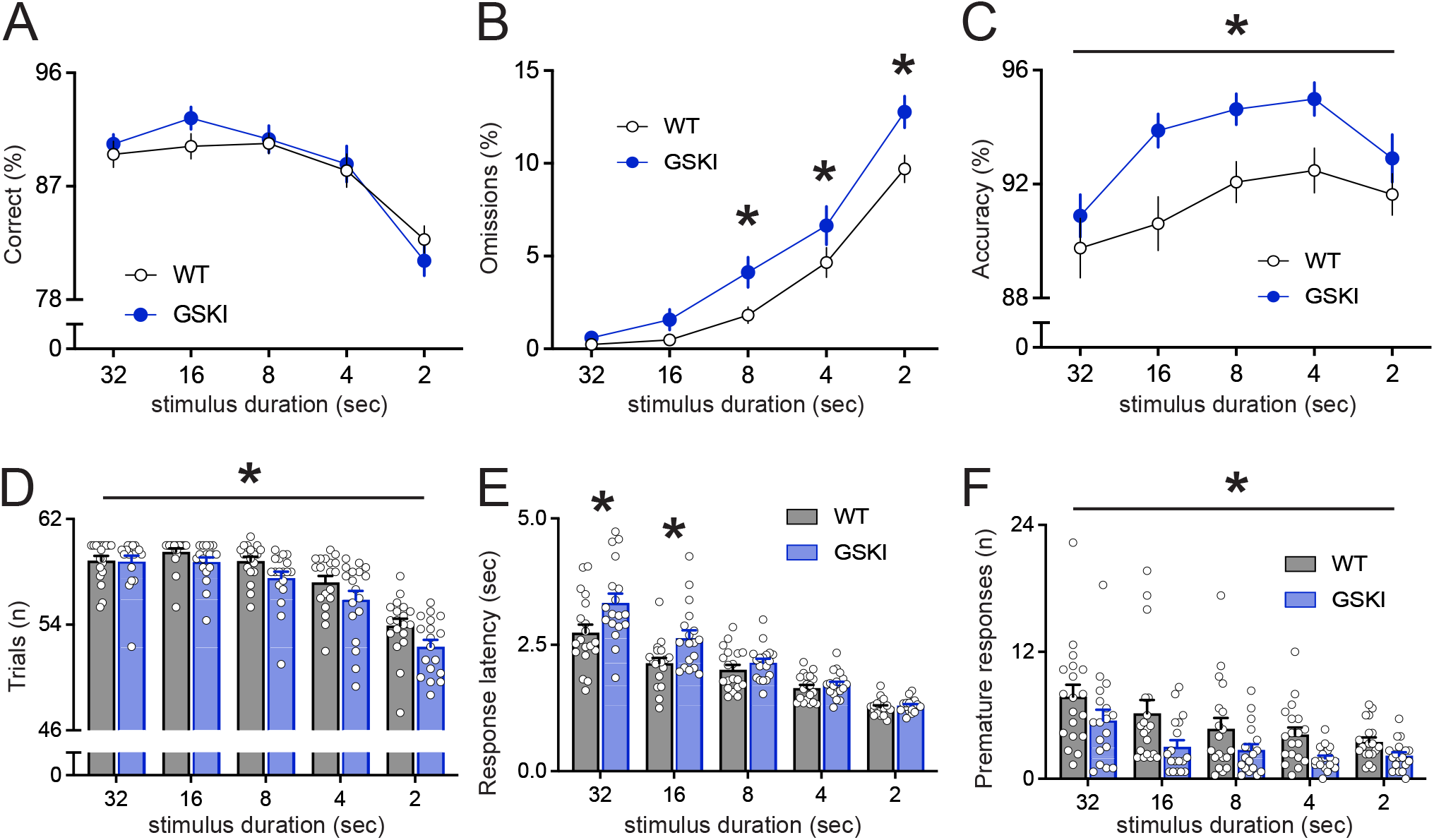
Training performance of WT and GSKI mice in the 5-CSRT task. **A**, WT (black lines) and GSKI (blue lines) showed similar percentages of correct responses across all stimulus durations [main effect (genotype): F_1,35_ = 0.195, p = 0.66]. **B**, GSKI mice made significantly more omissions [main effect (genotype): F_1,35_ = 11.26, p = 0.0019; genotype x stimulus duration, F_4,510_ = 2.72, p = 0.029; comparisons of estimated marginal means, 8 sec, p = .0035; 4 sec, p = 0.012; 2 sec, p = 0.0001]. **C**, GSKI mice showed significantly greater accuracy in comparison with WT mice [main effect (genotype): F_1,35_ = 8.62, p = 0.005, but no genotype x stimulus duration interaction, F_4,510_ = 1.58, p = 0.18]. **D**, GSKI mice (blue bars) completed fewer trials compared to WT mice (grey bars) [main effect (genotype): F_1,35_ = 5.78, p = 0.022, but no genotype x stimulus duration interaction, F_4,510_ = 1.92, p = 0.11. **E,** GSKI mice showed significantly longer correct response latencies in comparison with WT mice [main effect (genotype): F_1,35_ = 5.94, p = 0.02; genotype x stimulus duration interaction, F_4,510_ = 7.43, p < .0001; comparisons of estimated marginal means, 32 sec, p < 0.0001; 16 sec, p = 0.0005]. **F,** GSKI mice made significantly fewer premature responses in comparison with WT mice [main effect (genotype): F_1,35_ = 8.38, p = 0.0065, but no genotype x stimulus duration interaction, F_4,510_ = 0.67, p = 0.61]. n= 19 WT, 18 GSKI mice.

Once all mice achieved stable performance at baseline (2 sec stimulus duration), cognitive abilities were challenged in two probe trials. Poorer performance by the GSKI mice at the shortest stimulus durations also suggested attentional deficits, even after they had reached criterion on the basic task. In probe 1 (between-session design), entire sessions with stimulus durations of 1.5, 1.0, 0.8, 0.6 and 0.4 sec were presented in a random session order to prevent order effects. Mice were tested on a single stimulus duration for two consecutive days, interleaved with testing for two consecutive days at baseline (2 sec stimulus duration). In probe 1, GSKI mice showed a significantly greater percentage of omissions (**Fig. 2A**), a significantly lower percentage of correct responses (**Fig. 2B**) and completed significantly fewer trials (**Fig. 2D**) compared to WT mice. GSKI mice showed slightly worse response accuracy at the shortest stimulus duration (**Fig. 2C**). Additionally, GSKI mice showed a longer correct response latency at 0.6 sec and 0.4 sec (**Fig. 2E**). Moreover, there was a main effect of genotype on premature responding, with GSKI mice making significantly fewer premature responses than WT mice (**Fig. 2F**). There were no significant genotype differences in perseverative responses at any stimulus duration (genotype, F_1,~37.4_ = 1.06, p = 0.31, stimulus duration x genotype interaction: F_5,~676.4_ = 0.5, p = 0.77).

**Figure 2.**
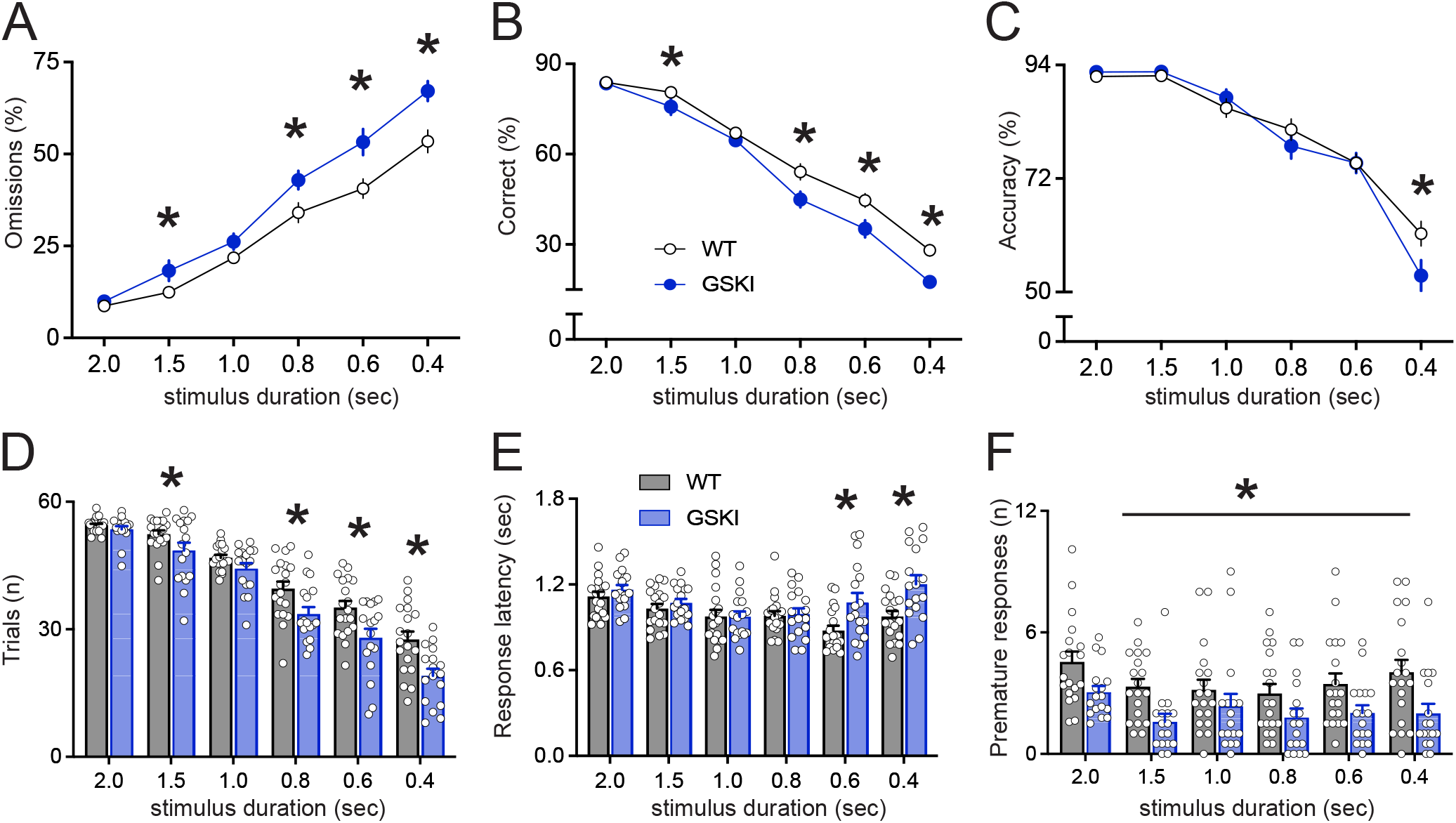
Performance of WT and GSKI mice on probe trial 1 in the 5-CSRT task. **A,** GSKI mice (blue lines) showed a greater percentage of omissions in comparison with WT mice (black lines) [main effect (genotype): F_1,~42.9_ = 36.3, p < 0.0001, stimulus duration x genotype: F_5,~676.7_ = 9.72, p < 0.0001; comparisons of estimated marginal means, 1.5 sec, p = .014; 0.8 sec, p = 0.0002; 0.6 and 0.4 sec, p < 0.0001]. **B,** GSKI mice showed a decrease in percentage of correct responses at the shortest stimulus duration [main effect (genotype): F_1,~40.1_ = 17.4, p = 0.0002, stimulus duration x genotype: F_5,~676.4_ = 8.06, p < 0.0001; comparisons of estimated marginal means, 1.5 sec, p = 0.047; 0.8 sec, p = 0.0002; 0.6 sec, p = 0.0001, 0.4 sec, p < 0.0001]. **C**, WT and GSKI showed similar accuracy across all stimulus durations except the shortest (0.4 sec) [main effect (genotype): F_1,~42.5_ = 1.33, p = 0.25, stimulus duration x genotype: F_5,~677_ = 5.59, p < 0.0001; comparisons of estimated marginal means, 0.4 sec, p < 0.0001]. **D**, GSKI mice (blue bars) completed fewer trials in comparison with WT mice (grey bars) [main effect (genotype): F_1,~41.7_ = 31.7, p < 0.0001, stimulus duration x genotype: F_5,~676.7_ = 8.94, p < 0.0001; comparisons of estimated marginal means, 1.5 sec, p = 0.01; 0.8 sec, p = 0.0001; 0.6 and 0.4 sec, p < 0.0001]. **E**, GSKI mice showed significantly longer correct response latencies at shorter stimulus durations [main effect (genotype): F_1,~39.3_ = 5.37, p = 0.026, stimulus duration x genotype: F_5,~676.9_ = 4.70, p = 0.0003; comparisons of estimated marginal means, 0.6 sec, p = 0.0008; 0.4 sec, p = 0.0001]. **F,** GSKI mice made significantly fewer premature responses during the ITI [main effect (genotype): F_1,~44.5_ = 8.99, p = 0.0044]. n = 19 WT, 17 GSKI mice.

In probe 2 (within-session design), stimulus durations of 2.0, 1.0, 0.8, and 0.6 sec were presented in random order within a session, with each stimulus duration presented an equal number of times per session. Similar to probe 1, GSKI mice displayed a significant increase in percent omissions (**Fig. 3A**) and a decrease in percent correct responses (**Fig. 3B**) at the shortest stimulus duration in comparison with WT mice. There was no significant difference in percent response accuracy (**Fig. 3C**), average correct response latencies (**Fig. 3D**), premature responses (**Fig. 3E**), or perseverative errors following correct responses (**Fig. 3F**) for the GSKI mice compared to WT mice.

**Figure 3.**
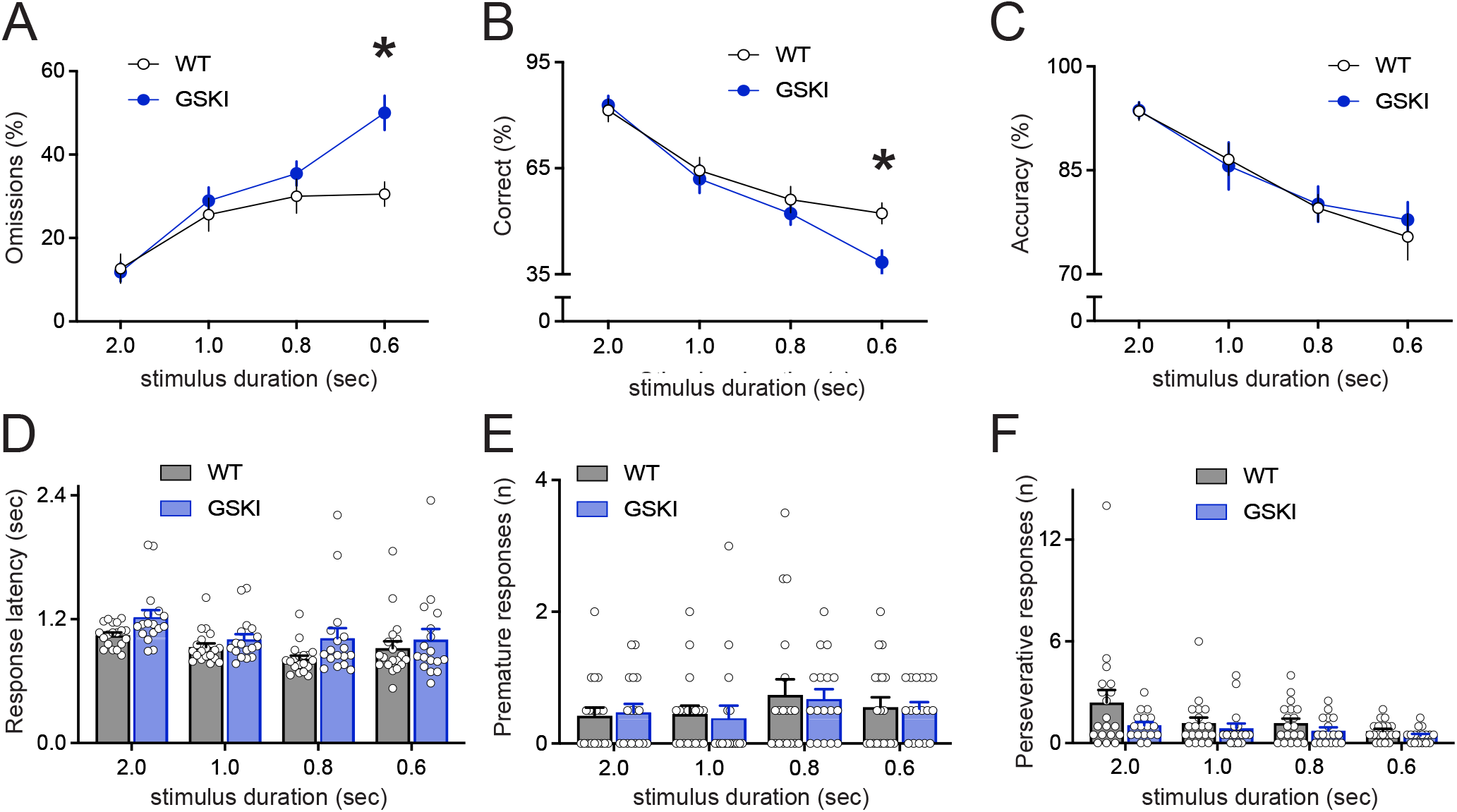
Performance of WT and GSKI mice on probe trial 2 in the 5-CSRT task. **A,** GSKI mice (blue line) displayed significantly more omissions at the shortest stimulus duration in comparison with WT mice (black line) [stimulus duration x genotype: F_3,246_ = 8.47, p < 0.0001; comparisons of estimated marginal means, 0.6 sec, p = 0.0001]. **B,** GSKI mice showed a decreased percentage of correct responses at the shortest stimulus duration [stimulus duration x genotype: F_3,246_ = 4.50, p = 0.0043; comparisons of estimated marginal means, 0.6 sec, p = 0.0001]. **C,** GSKI and WT mice showed similar accuracy [main effect(genotype): F_1,34_ = 0.052, p = 0.82; stimulus duration x genotype: F_3,246_ = 0.25, p = 0.86]. **D,** GSKI mice (blue bars) showed similar correct response latencies in comparison with WT mice (grey bars) [main effect (genotype): F_1,34_ = 3.38, p = 0.075; stimulus duration x genotype: F_3,246_ = 0.73, p = 0.54]. **E,** There were no significant differences between genotypes in premature responses [main effect (genotype): F_1,34_ = 0.024, p = 0.88; stimulus duration x genotype: F_3,246_ = 0.066, p = 0.98] nor in (**F**) perseverative responses [main effect (genotype): F_1,34_ = 3.88, p = 0.057; stimulus duration x genotype: F_3,246_ = 1.20, p = 0.31]. n= 13 WT, 11 GSKI mice.

### Deficits in 5-CSRT task not attributable to diminished motivation, locomotion or impaired visual sensory perception

The performance differences between GSKI and WT mice at the shortest stimulus durations could reflect deficits in divided attention or, alternatively, could reflect diminished motivation, locomotor deficits or impaired visual sensory perception. To investigate these alternative possibilities, upon completion of the 5-CSRT task, motivation was tested by subjecting a subset of mice to a progressive ratio (PR) task (Heath et al., 2016). Here, we found that neither the breakpoints (**Fig. 4A**) nor reward collection latencies (**Fig. 4B**) differed significantly between genotypes. Consistent with this, we found no differences between genotypes in reward collection latencies in either probe 1 (**Fig. 4C**) or probe 2 (**Fig. 4D**) in the 5-CSRT task. Thus, performance deficits of the GSKI mice are not likely attributable to diminished motivation.

**Figure 4.**
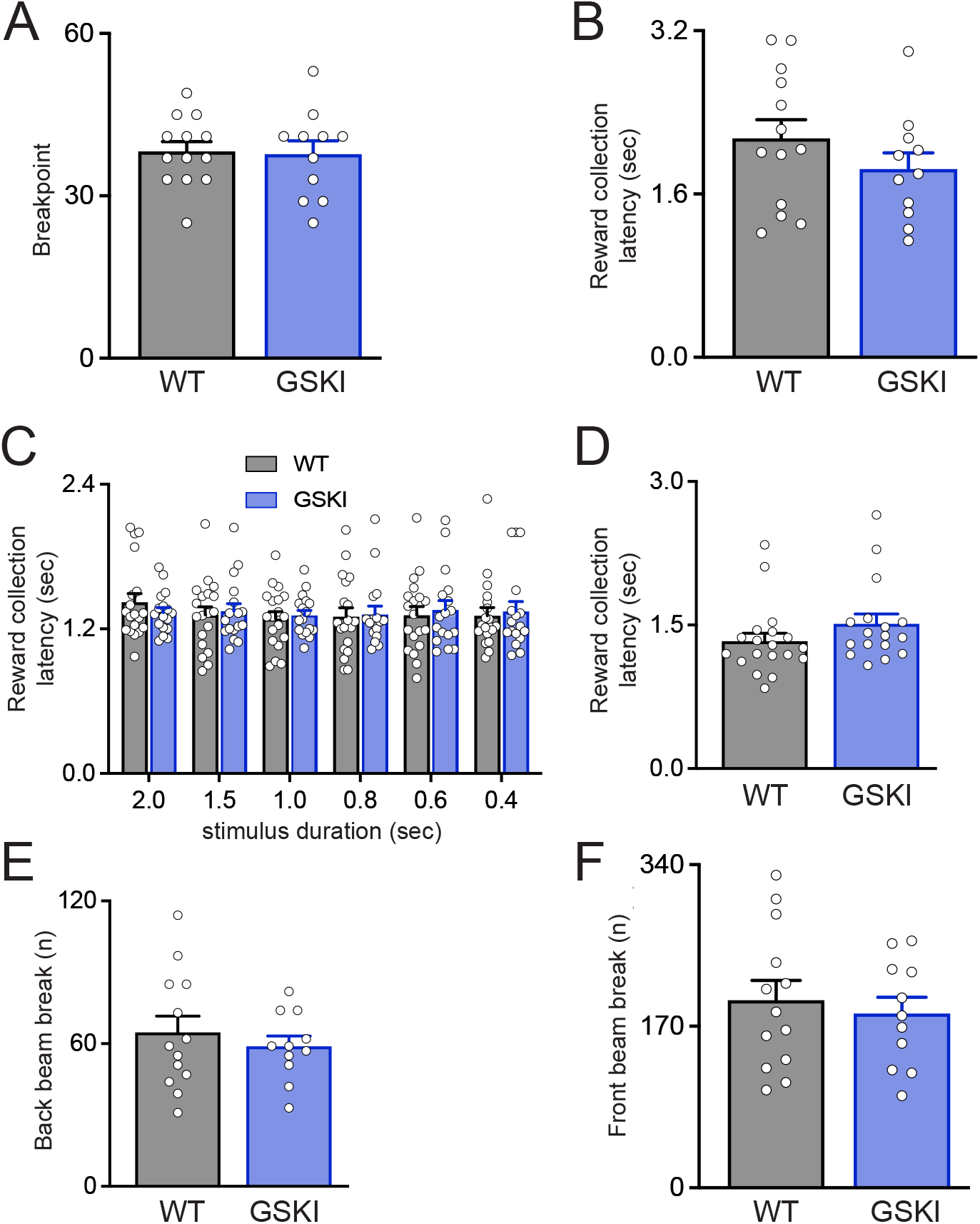
No differences between genotypes in motivation for reward-related behaviors or locomotor activity. In a progressive ratio (PR4) task to test for motivation, the breakpoints (**A**) and reward collection latencies (**B**) were similar between genotypes (breakpoints: Welch’s t-test, t_18.87_ = 0.166, p = 0.87; latencies: Welch’s t-test, t_21.93_ = 1.23, p = 0.23). Similarly, there were no differences between genotypes in reward collection latencies in probe 1 [**C,** main effect (genotype): F_1,~42.3_ = 0.046, p = 0.83] or probe 2 (**D,** Welch’s t-test, t_31.7_ = 1.37, P = 0.18) in the 5-CSRT task (n=13 WT, 11 GSKI mice) in the 5-CSRT. There were no differences between genotypes in numbers of back beam breaks (**E,** Welch’s t-test, t_19.73_ = 0.73, p = 0.48) or front beam breaks (**F,** Welch’s t-test, t_21.76_ = 0.53, p = 0.6) during the progressive ratio task, suggesting no differences in locomotion. For the progressive ratio task, n = 13 WT, 11 GSKI mice.

We next analyzed locomotor activity within the touch-screen chamber. There were no differences between genotypes in the number of back or front beam-breaks (**Figs. 4E, F**), suggesting no overt locomotor abnormalities. This conclusion is consistent with prior studies showing that locomotor activity of the GSKI mice during exploration of an arena is similar to that of WT mice (Yue et al., 2015; Matikainen-Ankney et al., 2018; Guevara et al., 2020).

Finally, to test visual sensory perception at the shortest stimulus durations of 0.6 and 0.4 sec, where deficits in the 5-CSRT task were manifest, we implemented a 1-choice version of the task by fixing the stimulus location to the middle response window only, thereby eliminating the requirement that mice divide their attention across 5 spatial locations (Humby et al., 1999). Under the lower attentional demands of this task, GSKI and WT mice showed no significant differences in omissions (**Fig. 5A**) or in the number of completed trials (**Fig. 5B**). These data suggest that visual sensory deficits per se are unlikely to underlie the higher percentage of omissions observed in the probe trials. However, there remained a main effect of genotype on correct response latency, with GSKI mice displaying significantly longer latencies (**Fig. 5C**). Thus, in the 1-choice task, omissions were normalized in the GSKI mice despite persistently slower information processing speed. This suggests that longer response latencies are unlikely to solely underlie the higher rate of omissions observed in the probe trials, though it may exacerbate the burden placed on attentional mechanisms when dividing attention across five screens.

**Figure 5.**
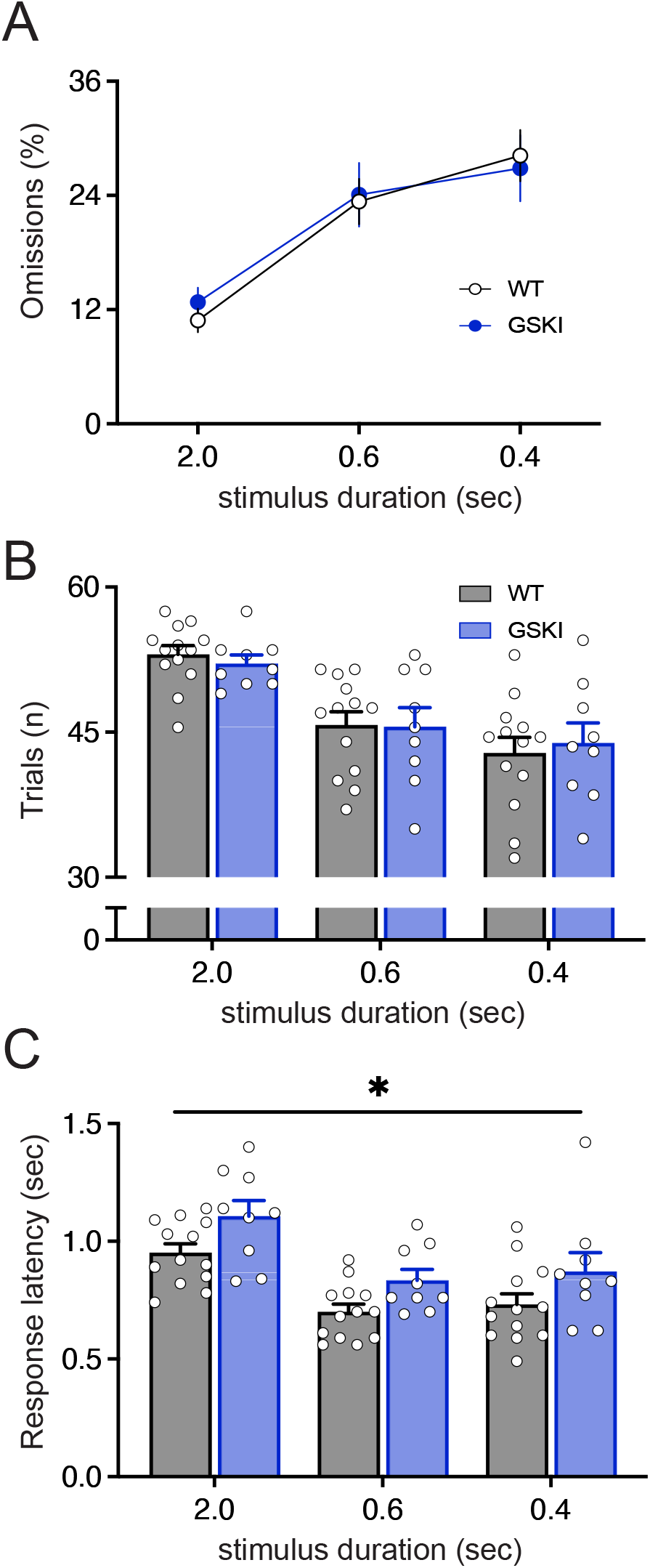
Reducing attentional demand in a 1-choice version of the serial reaction time task normalizes performance deficits in GSKI mice observed at the shortest stimulus durations. There were no differences between genotypes in percent omissions [main effect (genotype): F_1,20_ = 0.0304, p = 0.86; stimulus duration x genotype: F_2,106_ = 0.40, p = 0.67] or in number of completed trials [main effect (genotype): F_1,20_ = 0.0002, p = 0.99 stimulus duration x genotype: F_2,106_ = 0.35, p = 0.70] in the 1-choice task. There remained a main effect of genotype on correct response latency (**C**) similar to the 5-CSRT, with GSKI mice showing significantly longer response latencies [main effect (genotype): F_1,20_ = 5.24, p = 0.033]. n = 13 WT, 9 GSKI mice.

### Cholinergic signaling implicated in attention and processing speed deficits in the GSKI mice

Attentional processes are strongly modulated by central cholinergic mechanisms in rodents and humans (Baxter and Chiba, 1999). We therefore tested whether peripheral administration of the centrally-acting acetylcholinesterase inhibitor donepezil (Kosasa et al., 1999) would mitigate performance deficits displayed by the GSKI mice in the 5-CSRT task. Using a within-subject design, WT and GSKI mice received either saline or donepezil (0.3 mg/kg, IP), counterbalanced across mice of each genotype, one hour before testing with a 0.4 sec stimulus duration. We found that GSKI mice treated with donepezil made significantly fewer omissions (**Fig. 6A**) and completed more trials (**Fig. 6B**) than when treated with saline. Additionally, GSKI mice treated with donepezil showed shorter correct-response latencies in comparison with their performance when treated with saline (**Fig. 6C**). The GSKI mice displayed no significant differences in numbers of correct responses when treated with donepezil compared to when they were treated with saline (**Fig. 6D**). In contrast, in WT mice, there were no significant differences in any performance measure between saline and donepezil treatment (**Figs. 6E-H**). Together, these data suggest that acetylcholinesterase inhibition mitigated performance deficits displayed by the GSKI mice in the 5-CSRT task.

**Figure 6.**
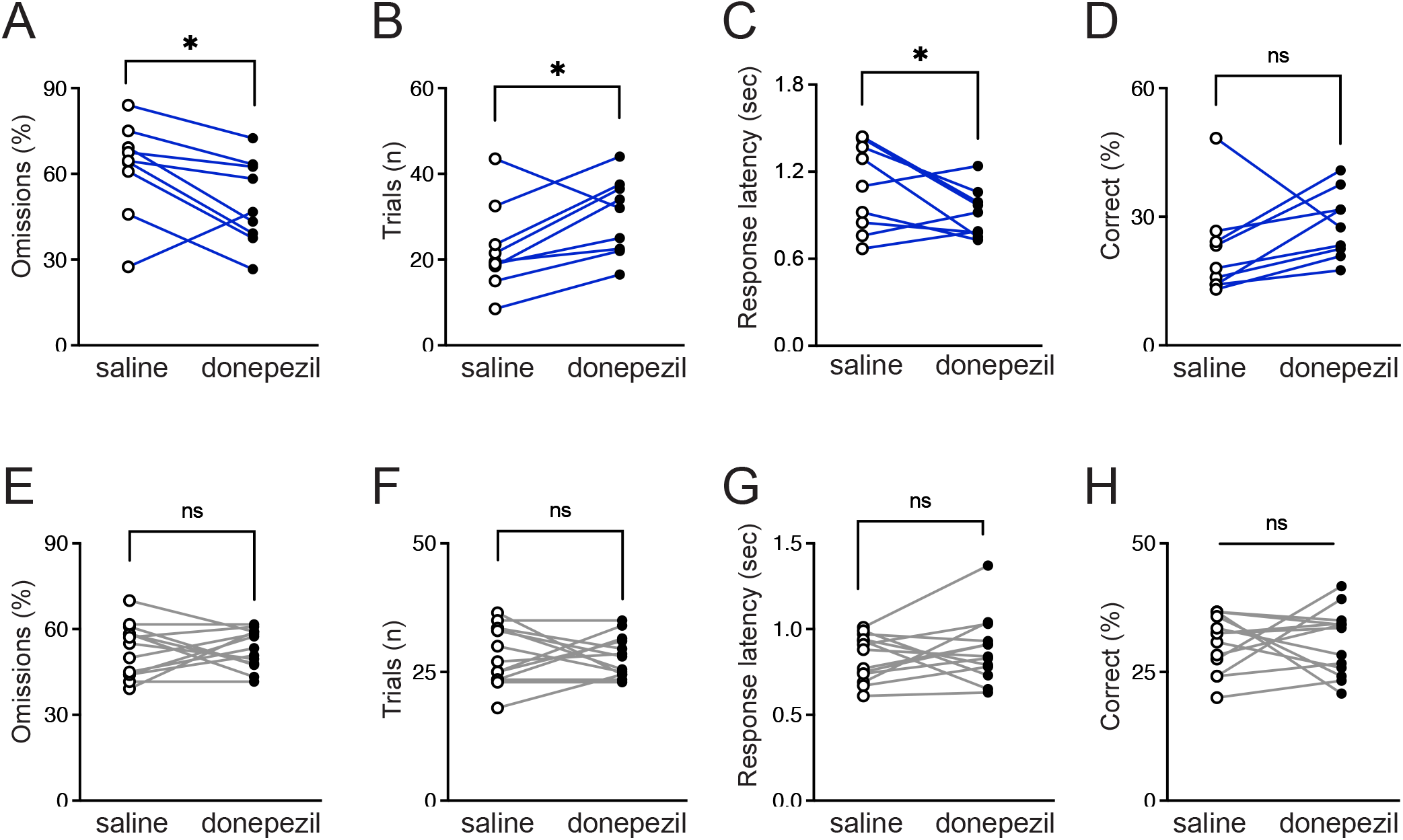
Systemic administration of donepezil, an acetylcholinesterase inhibitor, ameliorated performance deficits of the GSKI mice in the 5-CSRT task. Donepezil (0.3 mg/kg, IP) or saline was administered to GSKI mice (blue lines, **A-D**) or WT mice (grey lines, **E-H**), using a within-subject design, 1 hr before testing in the 5-CSRT task with a 0.4 sec stimulus duration. Compared to saline treatment (open circles), GSKI mice treated with donepezil (black circles) made significantly fewer omissions (**A,** genotype x treatment: F_1,64_ = 5.72, p = 0.020; comparisons of estimated marginal means, GSKI saline vs donepezil, p = 0.003; **E**, WT saline vs donepezil, p = 0.97) and showed an increased number of completed trials (**B,** genotype x treatment: F_1,64_ = 6.47, p = 0.013; comparisons of estimated marginal means, GSKI saline vs donepezil, p = 0.0016; **F**, WT saline vs donepezil, p = 0.98). Treating GSKI mice with donepezil lowered correct response latencies compared to saline treatment (**C,** genotype x treatment: F_1,64_ = 5.91, p = 0.018; comparisons of estimated marginal means, GSKI saline vs donepezil, p = 0.016; **F**, WT saline vs donepezil, p = 0.41). There was no effect of donepezil on percent correct responses (**D**, genotype x treatment: F_1,64_ = 2.15, p = 0.15). In contrast to GSKI mice, donepezil treatment in WT mice had no significant effects (ns) on omissions (**E**), number of completed trials (**F**), response latencies (**G**) or percent correct responses (**H**) in comparison with saline treatment. n = 13 WT, 9 GSKI mice.

### GSKI mice display altered cholinergic innervation of mPFC and striatum

To understand better a potential basis for the restorative effect of donepezil on attention and processing speed deficits in the young adult GSKI mice, we analyzed the density, size and intensity of immunofluorescent vesicular acetylcholine transporter (VAChT) labeling of cholinergic axons innervating mPFC (areas PL/IL) and dorsomedial striatum in GSKI and WT mice. In both structures, VAChT immunolabeling appeared as fragments of thin fibers studded with varicosities, similar to previous descriptions (Henny and Jones, 2008). We found significantly lower cholinergic fiber innervation densities and smaller immunofluorescent puncta sizes across all layers of mPFC (**Fig. 7A-C**) and in dorsomedial striatum (**Fig. 7D-F**) of GSKI mice in comparison with WT mice. There were no significant genotype differences in VAChT immunofluorescence intensity. These data reveal a significant anatomical alteration in cholinergic innervation of two principal interconnected structures that mediate executive function in young adult mutant mice, and suggest a basis for the rescue of some behavioral measures by donepezil.

**Figure 7.**
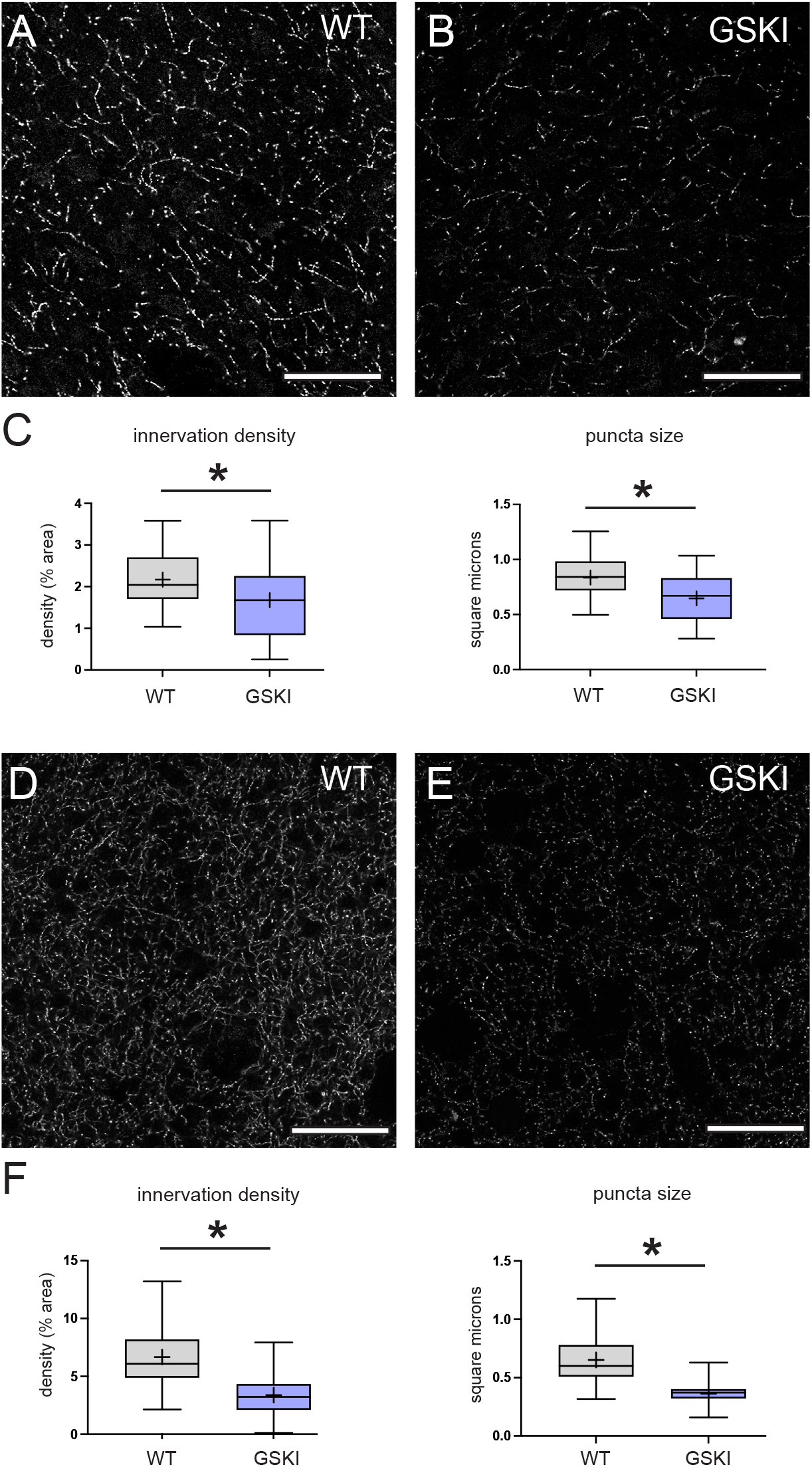
GSKI mice show altered pattern of cholinergic fiber innervation in mPFC and dorsomedial striatum in comparison with WT mice. Representative confocal microscope images of VAChT immunofluorescent labeling of cholinergic fibers in L5 of mPFC areas PL/IL (**A, B**) or dorsomedial striatum (**D, E**) in WT (left images of pair) or GSKI mice (right images of pair), as indicated. bars = 50 μm. **C, F,** Box-plots show that in PL/IL (**C**) and in dorsomedial striatum (**F**), fiber innervation density is significantly sparser and VAChT-labeled puncta size is significantly smaller in GSKI mice in comparison with WT mice (fiber innervation density, main effect of genotype, F_1,~4.2_ = 9.24, p = 0.036; puncta size, main effect of genotype, F_1,~4.2_ = 9.24, p = 0.043). n = 3 WT, 3 GSKI mice.

### GSKI mice are impaired in goal-directed learning

We assayed goal-directed actions using a 4-day instrumental conditioning paradigm that in WT mice produces goal-directed action-outcome associations verified subsequently by sensitivity to outcome devaluation (Shan et al., 2014). Both WT and GSKI mice made a similar number of responses under a continuous reinforcement (CRF) schedule on day 1 (**Fig. 8A**). On subsequent days 2-4 of the instrumental task, the reward was delivered on a random interval (RI) schedule of 15, 30 and 30 sec. As expected, WT mice made progressively greater numbers of responses on days 2-4 (**Fig. 8A**) and, following the 4-day instrumental conditioning paradigm, they exhibited sensitivity to reward devaluation (**Fig. 8B**), indicating goal-directed learning as expected (Shan et al., 2014). In contrast, GSKI mice made significantly fewer responses to the stimulus under the RI schedules on days 3 and 4 in comparison with WT mice (**Fig. 8A**), and were insensitive to outcome devaluation (**Fig. 8B**), suggesting an impairment in instrumental conditioning and goal-directed learning.

**Figure 8.**
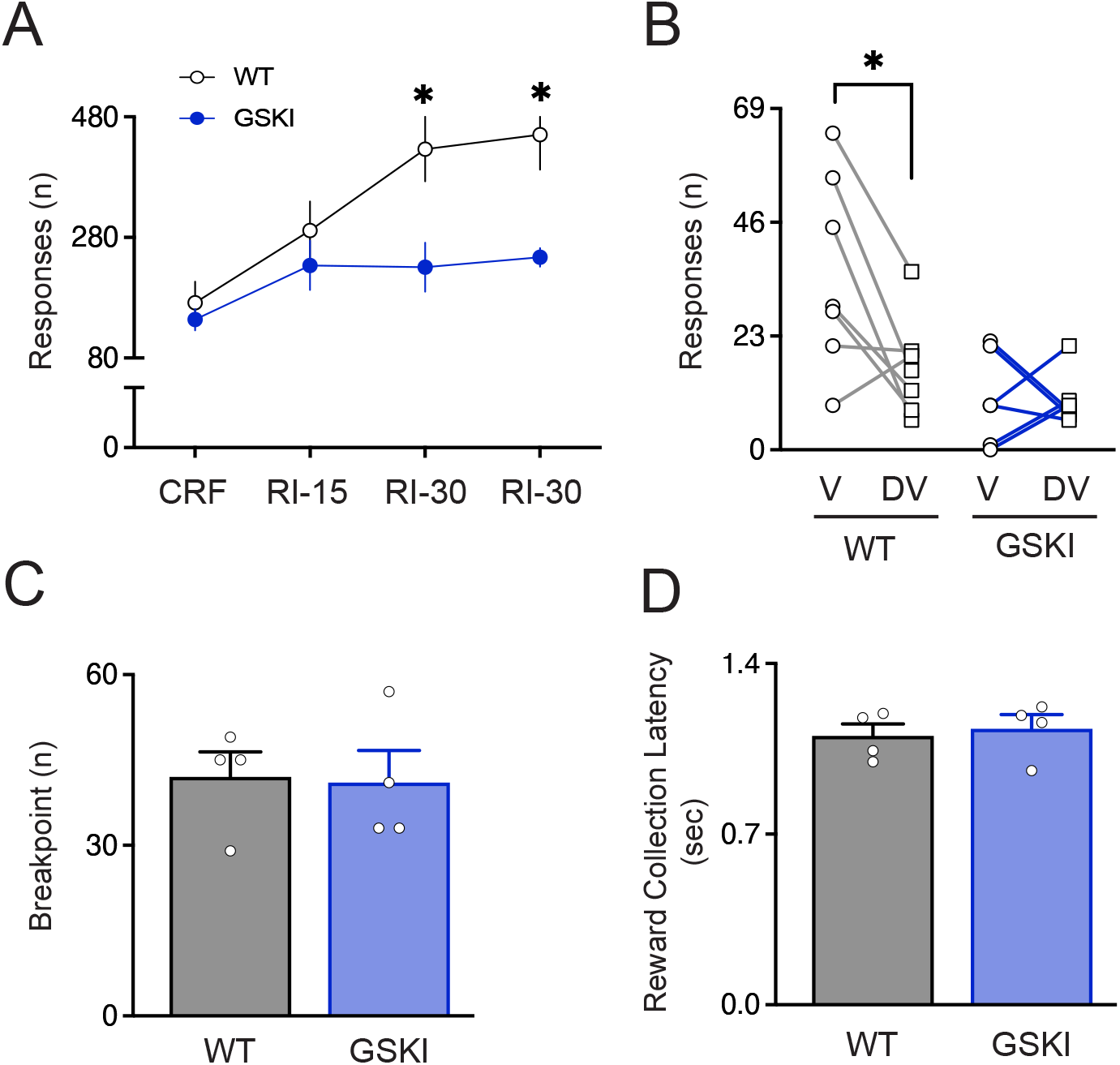
GSKI mice are impaired in goal-directed learning. WT and GSKI mice underwent a 4 day instrumental conditioning task followed by outcome devaluation to assay goal-directed learning. **A**, WT mice (black line) made significantly greater numbers of responses during the RI schedule (x-axis), while GSKI mice (blue line) made significantly fewer responses compared to WT on days 3 and 4 RI schedules [main effect (genotype), F_1,13_ = 5.77, p = 0.033, genotype x schedule: F_3,39_ = 5.92, p = 0.002; comparison of estimated marginal means, RI-30 day 3, p = 0.0034; RI-30 day 4, p = 0.0025]. CRF, continuous reinforcement; RI-15, random interval 15 sec; RI-30, random interval 30 sec. **B**, WT (grey lines, left) but not GSKI mice (blue lines, right) demonstrated goal-directed outcome association by their sensitivity to outcome devaluation (valued condition, V = chow; versus devalued condition, DV = strawberry milk) [main effect (genotype), F_1,11_ = 8.65, p = 0.013; genotype x condition: F_1,11_ = 4.641, p =0.054; comparison of estimated marginal means, WT chow vs SM p = 0.0085, GSKI chow vs SM p = 0.98]. n =7 WT, 6 GSKI mice for data shown in **A**, **B**. In subsequent tests of motivation for the reward, neither breakpoints in a PR4 task (**C**) nor reward collection latencies (**D**) differed between genotypes (breakpoints: Welch’s t-test, t_5.68_ = 0.14, p = 0.89; latencies: Welch’s t-test, t_5.83_ = 0.38, p = 0.72). n = 4 WT, 4 GSKI mice for data shown in **C, D**.

Goal-directed action-outcome association is also sensitive to motivation (Heath et al., 2016). Thus, following the last outcome devaluation test, we subjected the WT and GSKI mice to a PR task to compare their motivation for the reward. As before, we found no significant differences between genotypes in break points (**Fig. 8C**). Additionally, there were no differences between genotypes in average reward collection latencies (**Fig. 8D**). These data suggest that alterations in motivation for the reward are unlikely to underlie deficits in goal-directed learning in GSKI mice.

### Cognitive flexibility is unaffected in GSKI mice

We assayed cognitive flexibility with a pairwise visual discrimination task followed by reversal learning. During the initial visual discrimination phase, we found no significant differences between genotypes in the number of sessions required to reach performance criteria or in percent correct responses (**Fig. 9A**), in session length (**Fig. 9B**) or in correction responses (**Fig. 9C**).

**Figure 9.**
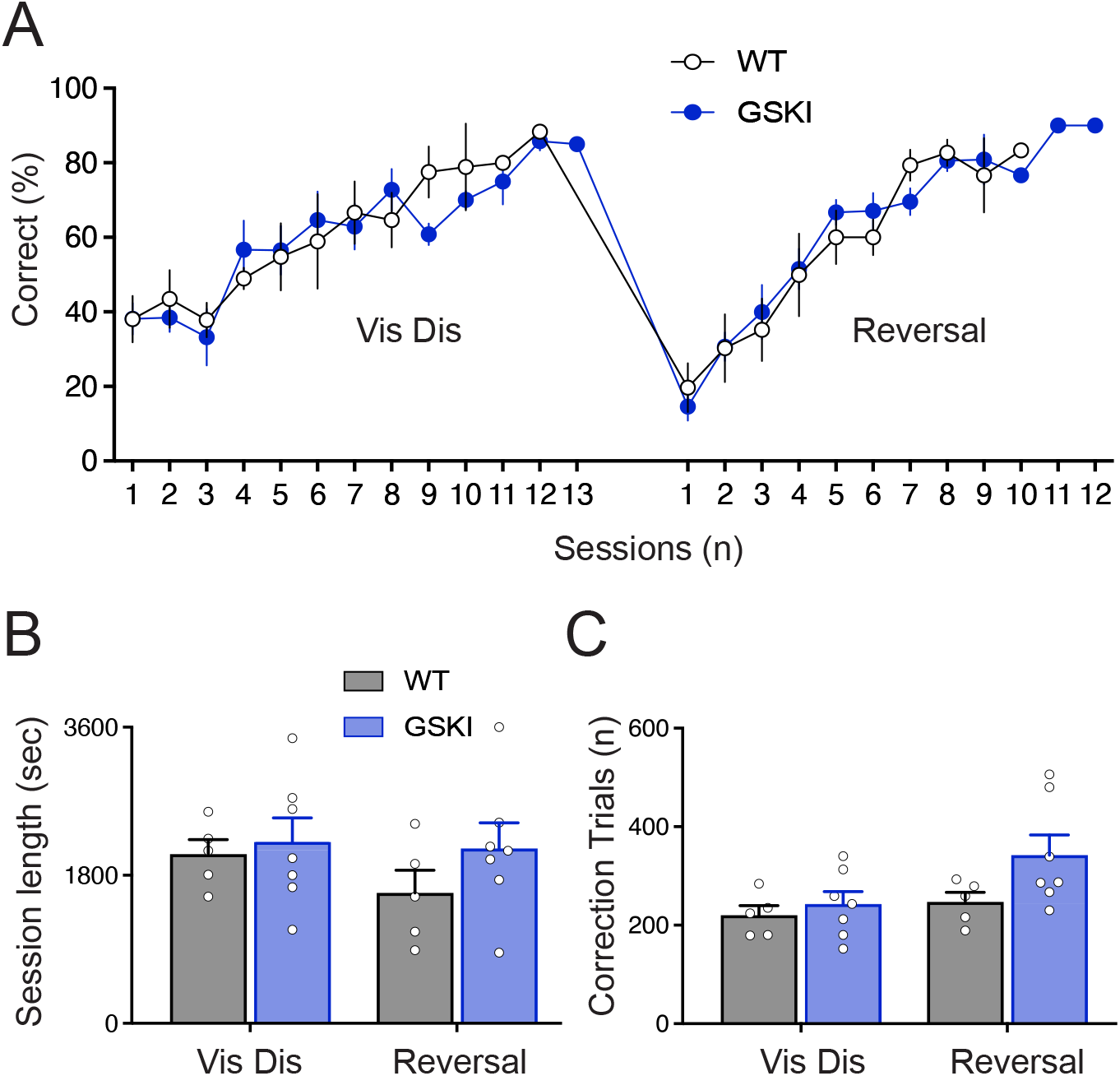
Pairwise visual discrimination and reversal learning, a measure of cognitive flexibility, is not significantly different in GSKI mice in comparison with WT mice. **A**, GSKI mice reached performance criteria during both the visual discrimination phase (Vis Dis) and subsequent reversal learning phase (Reversal) over a similar number of sessions in comparison with WT mice (main effect of genotype and interactions of genotype with session and testing phase, Fs < 1, ps > 0.35). **B,** there were no genotype differences observed in session length during visual discrimination (Vis Dis) (main effect of genotype F_1,~23.1_ = .0048, p = 0.95; genotype x testing phase, F_1,~213.1_ = .1.57, p = 0.21)]. **C,** Mice made more correction trials overall in the reversal phase compared to the initial discrimination phase, F_1,~202.1_ = 73.3, p < 0.0001, but there was no main effect of genotype on correction responses, F_1,~53.7_ = .035, p = 0.85, or genotype x phase interaction, F_1,~202.1_ = 2.77, p = 0.098, suggesting stimulus-reward learning is intact in GSKI mice. n = 5 WT, 7 GSKI mice.

In the subsequent reversal phase, when the stimulus/reward contingency was reversed, GSKI and WT mice reached performance criteria after a similar number of sessions, with similar percentages of correct responses (**Fig. 9A**) and session lengths (**Fig. 9B**). WT mice showed no significant difference in the number of corrective responses between the discrimination and reversal phases (**Fig. 9C**), and while GSKI mice showed a slightly greater number of correction responses in the reversal phase in comparison with their number of correction responses in the discrimination phase, there were no significant genotype x phase interactions (**Fig. 9C**), suggesting that stimulusreward learning was intact in the GSKI mice.

## DISCUSSION

We used visuospatial touchscreen-based operant tasks to assess executive function in young adult male mice carrying a PD-associated *Lrrk2*-G2019S knockin mutation. GSKI mice displayed significant deficits in visuospatial attention and longer response latencies, suggesting significantly slower information processing. GSKI mice were also impaired in instrumental conditioning and goal-directed learning, while pairwise visual discrimination and reversal learning--a measure of cognitive flexibility--appeared largely unaffected. In humans, deficits in attention and processing speed are characteristics of early cognitive symptoms of idiopathic and several familial forms of PD, including asymptomatic *LRRK2*-G2019S carriers (Thaler et al., 2012; Robbins and Cools, 2014; Perugini et al., 2018; Piredda et al., 2020). We found that measures of impaired attention in the GSKI mice were rescued by donepezil, a centrally-acting acetylcholinesterase inhibitor, and that cholinergic innervation of mPFC and striatum was significantly sparser than WT, suggesting that altered cholinergic anatomy in brain may have direct or indirect contributions to cognitive dysfunction in young adult GSKI mice. Taken together, these data suggest a framework for better understanding deficient cognitive control mechanisms that may underlie early PD-associated nonmotor cognitive impairment.

The 5-CSRT task is a sensitive test of visuospatial attention in rodents (Robbins, 2002; Amitai and Markou, 2010). In the GSKI mice, the main hallmark of an attention deficit was a consistently higher percentage of omissions, evident during the training phase but significantly affecting performance at the shortest stimulus durations in the two probe trials when attentional demand is greatest. Control experiments ruled out motivational, locomotor or visual sensory perceptual deficits, thus suggesting a general deficit in divided attention in the GSKI mice. The significantly longer response latencies in probe 1 suggest that GSKI mice have slower information/decision processing speed compared to WT mice, which likely contributed to the significantly greater percentage of omissions. In contrast, response accuracy in the GSKI mice was not significantly different than WT mice. The combination of slower processing speed but preserved accuracy in the mutant mice is a performance profile similar to that described in a study of human non-PD manifesting carriers of the *LRRK2*-G2019S mutation on the Stroop Color-Word test, where significant performance differences were found in comparison with healthy controls (Thaler et al., 2012). Additionally, although the underlying circuitry involved in such deficits in the mice is currently unknown, generally each of the tasks that we used test cognitive domains that require intact connectivity between areas of medial and orbital prefrontal cortex and dorsomedial striatum (Chudasama and Robbins, 2004; Yin et al., 2005; Izquierdo et al., 2017). In human non-PD manifesting *LRRK2*-G2019S carriers, fMRI imaging during performance in the Stroop Color-Word test indicated abnormal connectivity between frontal and parietal cortical regions and striatum (Thaler et al., 2013; Bregman et al., 2017), which together suggest some similarities in neural mechanisms and circuits underlying aspects of executive dysfunction in mice and humans.

Some of the various behavioral deficits in the GSKI mice may reflect dysfunction in action selection, which could include faulty behavioral inhibition and/or faulty selection of an appropriate response (Aron, 2007; Friedman and Robbins, 2021). In addition to omissions, GSKI mice made significantly fewer premature responses in the 5-CSRT task (a heightened withholding of responses). While the precise neural basis for such deficits is unclear, our finding that pretreating GSKI mice with the acetylcholinesterase inhibitor donepezil--which increases the perisynaptic concentration of acetylcholine (Kosasa et al., 1999)--reversed the high rate of omissions and largely normalized response latencies in the 5-CSRT task implicates cholinergic signaling. In human idiopathic PD, cholinergic function in brain is reduced in both non-demented and demented patients, affecting cognition and other non-motor symptoms (Perez-Lloret and Barrantes, 2016). Similarly, PET studies of human *LRRK2*-mutation carriers, including those with the G2019S mutation, suggest that *LRRK2* mutation carriers with or without PD exhibit heightened acetylcholinesterase activity in several cortical and subcortical regions in comparison with idiopathic PD or healthy controls (Liu et al., 2018). This suggests that deficient cholinergic modulation of circuit function can be evident early, before motor symptoms, which could contribute to early cognitive dysfunction. On the other hand, while acetylcholinesterase inhibitors, including donepezil, have been used clinically to ameliorate cognitive decline in demented PD and Alzheimer’s patients (Simuni et al., 2009; Sharma, 2019; Armstrong and Okun, 2020), their effectiveness for improving mild cognitive impairment in early-stage PD is equivocal (Mamikonyan et al., 2015; Baik et al., 2021).

We found no effect of donepezil in WT mice, making it unlikely that the drug non-specifically elevated motivation or arousal. Rather, acetylcholine signaling, particularly in mPFC, modulates visuospatial attention specifically (Baxter and Chiba, 1999; Bloem et al., 2014). Acetylcholine transients across multiple time-scales are detected within mPFC of rodents during performance of visuospatial tasks that place demands on attentional mechanisms (Arnold et al., 2002; Parikh et al., 2007), including the 5-CSRT task (Passetti et al., 2000), and are triggered by cognitive operations that govern the transition from cue detection monitoring to activation of cue-directed responses (Parikh et al., 2007; Howe et al., 2013; Sarter et al., 2014). Such cue-associated acetylcholine transients reflect innervation of mPFC by basal forebrain cholinergic neurons (Ballinger et al., 2016), but extrinsic and intrinsic cholinergic innervation of striatum is also relevant to attentional and other cognitive control mechanisms. Lesions of striatally-projecting brainstem cholinergic neurons impair attention and reduce response latency in a 5-CSRT task (Cyr et al., 2015), while lesions of striatal cholinergic interneurons impair cognitive flexibility (Ragozzino et al., 2009; Bradfield et al., 2013; Aoki et al., 2015). One limit of our study is that donepezil was administered systemically, so it is unknown which cholinergic system is involved. Future studies will be needed to address this.

Although the reversal of some aberrant behavioral measures by donepezil in the 5-CSRT task implicates cholinergic signaling, the mechanism that links LRRK2 mutation to cholinergic function, either directly or indirectly, remains speculative. Our analyses of cholinergic innervation of mPFC and dorsomedial striatum indicate that in both structures, cholinergic innervation is significantly less dense in mutants, with smaller immunofluorescently-labeled puncta. Whether lower fiber density reflects fewer cholinergic somata or preserved somata each with sparser terminal axonal fields remains to be determined. The presence of overt anatomical abnormalities in VAChT labeling in young adults is striking in that striatal tyrosine hydroxylase levels and labeling appear normal at similar ages in G2019S knockin mice, although striatal dopamine signaling may be altered early (Volta et al., 2017). Cholinergic neurons may be precociously vulnerable to PD pathogenesis compared to DA neurons, perhaps explaining cognitive abnormalities that often precede abundant DA neuron degeneration. Mechanistically, early vulnerability could reflect G2019S-mediated disruption of trophic support of cholinergic neurons. Previous studies have shown that pathogenic LRRK2 mutations disrupt ciliogenesis in cortical and striatal neurons (Dhekne et al., 2018; Lara Ordónez et al., 2019; Khan et al., 2021), interfering with a cilia-dependent, reciprocal molecular signaling loop that neurotrophically maintains the nigrostriatal circuit (Gonzalez-Reyes et al., 2012). In this signaling loop, striatal DA axons secrete Sonic hedgehog (Shh) that is sensed by cilia on striatal cholinergic interneurons. They, in turn, respond to Shh by secreting GDNF, providing trophic support of both DA and striatal cholinergic neurons. Developing and adult basal forebrain cholinergic neurons also synthesize Shh (Reilly et al., 2002) and require NGF--which in mPFC is secreted by interneurons (Biane et al., 2014)--for survival and maintenance of terminal arbors (Lad et al., 2003). *In vitro* studies show synergistic trophic support of basal forebrain cholinergic neurons in the presence of both Shh and NGF (Reilly et al., 2002). Thus, it is plausible that G2019S-mediated disruption of neuronal cilia in mPFC or striatum progressively disrupts a neurotrophic loop akin to that described above, derailing proper maintenance of cholinergic innervation and function.

LRRK2-G2019S can also alter properties of presynaptic neurotransmitter release (Beccano-Kelly et al., 2014; Cirnaru et al., 2014; Pan et al., 2017). While the mutation may affect release of acetylcholine, a direct effect is unlikely, since expression levels of LRRK2 are very low in basal forebrain, cholinergic brainstem nuclei and striatal cholinergic interneurons. In contrast, LRRK2 expression is high in cortical neurons and in striatal projection neurons (SPNs) (Taymans et al., 2006; Giesert et al., 2013; West et al., 2014; Gokce et al., 2016), suggesting that functionally abnormal presynaptic input to cholinergic cells from G2019S-expressing cortical neurons alters appropriate top-down regulation of cholinergic cell firing. A final possibility is that convergent cholinergic and glutamatergic inputs interact to mitigate G2019S-driven deficits in glutamatergic synaptic plasticity that have been described in G2019S knockin mice. The stoichiometry of AMPA-type glutamate receptor subunits at excitatory synapses onto SPNs in adult G2019S knockin mice is altered in comparison with WT striatal synapses (Matikainen-Ankney et al., 2018; Chen et al., 2020), and in these mice, both direct- and indirectpathway SPNs in dorsomedial striatum are unable to express corticostriatal long-term potentiation (Matikainen-Ankney et al., 2018). Normal bidirectional synaptic plasticity of corticostriatal synapses is associated with the cognitive processes tested here (Lovinger, 2010; Shan et al., 2014; Marquardt et al., 2021), and cholinergic signaling in mPFC, striatum and elsewhere critically regulates the expression of persistent forms of glutamatergic synaptic plasticity (Calabresi et al., 1999; Centonze et al., 2003; Gong and Ford, 2019; Fernández de Sevilla et al., 2021; Cools and Arnsten, 2022). It is therefore plausible that cognitive deficits in the mutants reflect such abnormal glutamatergic synaptic signaling and plasticity in dorsomedial striatum, and that boosting ambient acetylcholine levels by donepezil is sufficient to drive convergent intracellular signaling pathways to compensate for such plasticity deficits.

## Acknowledgements

This work was supported by the National Institutes of Health, NINDS grant numbers F31NS117089 and R01NS107512. We thank Lydia Gould and Pamela Del Valle for expert technical assistance. Microscopy was performed at the Microscopy Core and Advanced Bioimaging Center at the Icahn School of Medicine at Mount Sinai.

